# Mutation rates in seeds and seed-banking influence substitution rates across the angiosperm phylogeny

**DOI:** 10.1101/156398

**Authors:** Marcel Dann, Sidonie Bellot, Sylwia Schepella, Hanno Schaefer, Aurélien Tellier

## Abstract

**Background:** Seed-banking (the ability to persist in the soil over many generations) is usually considered as a dormant stage where genotypes are “stored” as a bet-hedging strategy in response to unpredictable environments. However, seed dormancy may instead have consequences for the integrity of the DNA and generate novel mutations.

**Methods:** We address this paradox by building phylogenies based on the plastomes and nuclear ITS of species belonging to ten angiosperm clades. In each clade, the substitution rate (branch-length) of a seed-banking species is compared with that of a closely-related non-seed-banking species.

**Results:** Seed-banking species show as high or higher substitution rates than non-seedbanking species, and therefore mutations occur in dormant seeds at a rate at least as high as in above-ground plants. Moreover, seed born mutations have the same probability to reach fixation as those from above ground. Our results are robust to differences in selection, generation time, and polymorphism.

**Conclusions:** Mutations occurring in seeds, and thus seed-banking, affect the population diversity of plant species, and are observable at the macro-evolutionary scale. Our study has consequences for seed storage projects, since the stored seeds are likely to accumulate mutations at a higher rate than previously thought.

## Introduction

Seed-banking or long term dormancy is a prevalent strategy in many plant species but also in bacteria and invertebrates. This bet-hedging strategy evolves to maximize the geometric fitness of the population under variable and unpredictable environmental conditions (Brown & Venable, 1986; Evans & Dennehy, 2005). Seed-banking has evolved multiple times in angiosperms (Willis et al., 2014), and is frequently seen as an adaptation to desert habitats (Brown & Venable, 1986; Pake & Venable, 1996). Multiple adaptations at the physiological level allow dormancy, such as low metabolism or thick pericarp (Finch-Savage & Leubner-Metzger, 2006). Different types of dormancy can be distinguished based on the physiology and triggers to lift up the dormant state (Willis et al., 2014), and result in different types of seed-banks depending on the maximal length of dormancy: transient (< one year), short term persistent (one to five years), and long term persistent (≥ five years) (Thompson et al., 1997; Baskin & Baskin, 2014).

Seed-banking generates a so-called storage effect of diversity in the soil which has known consequences at the population level such as increasing genetic diversity (Templeton & Levin, 1979; Nunney, 2002; Lundemo et al., 2009), slowing down natural selection (Hairston Jr & De Stasio Jr, 1988; Koopmann et al., 2016), promoting balanced polymorphism (Turelli et al., 2001; Tellier & Brown, 2009) and decreasing genetic differentiation among populations (Vitalis et al., 2004; Falahati-Anbaran et al., 2014). The storage effect has also the consequence to buffer population size changes (Nunney, 2002), and decrease population extinction rates, which is known as the “rescue effect” (Brown & Kodric-Brown, 1977). The chief effect of longer term seed-banking is thus to increase the observable genetic polymorphism within a population both in seeds and in above-ground plants, irrespective of the origin of the mutation (above-ground plants or seeds).

In addition to the storage effect, Levin (Levin, 1990) proposed that seed-banks would not only conserve alleles over generations, but could also be a source of new alleles arising from mutations accumulating during a long stay in the soil. This view is supported by physiological and molecular biology studies demonstrating that DNA degradation occurs in seeds (Abdalla & Roberts, 1969; Cheah & Osborne, 1978; Murata et al., 1982; Chauhan & Swaminathan, 1984; Dourado & Roberts, 1984a; Dourado & Roberts, 1984b; Dandoy et al., 1987). Repair mechanisms of DNA in seeds have been recently elucidated, linking the maintenance of genome stability with the progression through germination (Waterworth et al., 2016). It follows therefore that species with longer seed-banks are expected to exhibit a higher rate of neutral, deleterious and advantageous mutations in seeds (Levin, 1990). This seems to be confirmed by a study of 16 pairs of closely-related seed-banking and nonseed-banking angiosperm species where the seed-bankers were found to have higher substitution rates (Whittle, 2006). However, this preliminary work was limited to the < 500 bp nuclear ITS region and did not control for possible confounding factors. In contrast to those findings, ecologists and population geneticists have predominantly assumed that mutations occurring in seeds are chiefly deleterious, and that neutral or advantageous mutations occur only in above-ground plants (Kaj et al., 2001; Vitalis et al., 2004; Tellier et al., 2011; Koopmann et al., 2016). Indeed, a meta-analysis comparing genetic diversity in the soil seed-banks and above-ground plants has revealed only marginal differences (Honnay et al., 2008). This is not surprising considering the short time scales of the study, and that the above-ground population is in effect a sub-sample of the seed-bank (Lundemo et al., 2009). However, a slight excess of rare alleles could be observed in the seed-banks (Honnay et al., 2008) based on various markers which provide imprecise knowledge about the mutation rate (e.g. AFLP or microsatellites). This excess has been attributed to possible selection at the seedling stage against deleterious mutations arising in seeds (Vitalis et al., 2004). To resolve this contradictory evidence, we analyze large chloroplast DNA regions (>>10 kb) and the nuclear ITS of selected angiosperm species with different seed-bank lengths, and quantify the amount of new mutations arising in seeds that become fixed at the macro-evolutionary time scale.

Resolving this issue has theoretical and empirical importance. First, in models investigating the effect of seed-banks on neutral and selected diversity in plant genomes (Kaj et al., 2001; Vitalis et al., 2004; Tellier et al., 2011; Koopmann et al., 2016) the per site per generation mutation rate is a crucial parameter, influencing the inference of short time scale evolutionary parameters (Živković & Tellier, 2012). Second, at the phylogenetic time scale, substitution rates are scaled per generation which means that the length of seed-banking may factor indirectly in branch length (Charlesworth, 1994) under the assumption that mutations occurring in seeds may reach fixation. Differences in seed-banking would thus have to be considered when investigating the origin of substitution rate heterogeneity in species trees, and when deciding a priori between strict or relaxed molecular clock models to time-calibrate a phylogeny. The effect of seed-banking and the mutation rate in seeds has so far been ignored in phylogenetic studies. Third, knowing if mutations occur in seeds and at which rate is of high relevance to sustain seed storage of plant species for conservation purposes (e.g., Millenium Seed-bank, http://www.kew.org/scienceconservation/collections/millennium-seed-bank).

In this study, we investigate the long-term effect of seed mutations on species substitution rates. We aimed to compare the substitution rates of species producing only transient seed-banks, i.e. short or non-seed-banking (NSB) species, and species producing long-term persistent seed-banks, i.e. seed-banking (SB) species sensu Thompson et al. (1997). In order to avoid confounding effects, we follow a “species-pair” approach as advocated in Lanfear et al. (Lanfear et al., 2010) by comparing substitution rates of closely-related and ecologically similar SB and NSB species forming pairs scattered across the angiosperm phylogeny (see Fig. 1). We estimate substitution rates from branch lengths obtained from the Bayesian phylogenetic analysis of 41 full (chloroplast) plastomes, 27 of which are newly sequenced and assembled for this study, while 14 are recovered from databases. These plastomes are obtained by both genome skimming of DNA from old herbarium specimens and using a newly developed protocol for chloroplast DNA enrichment. We use three different statistical tests to demonstrate that seed-banking species have increased substitution rates compared to non-seed-banking species. Furthermore, we test that this effect is not confounded by generation time or selection pressure. In conclusion, we call for more ecological studies of the intensity and age of seed-banking for individual plant lineages, and we encourage population geneticists and phylogeneticists to pay more attention to seed mutations and differences in seed-banking when they elaborate models of micro- and macro-evolution.

**Figure 1.**
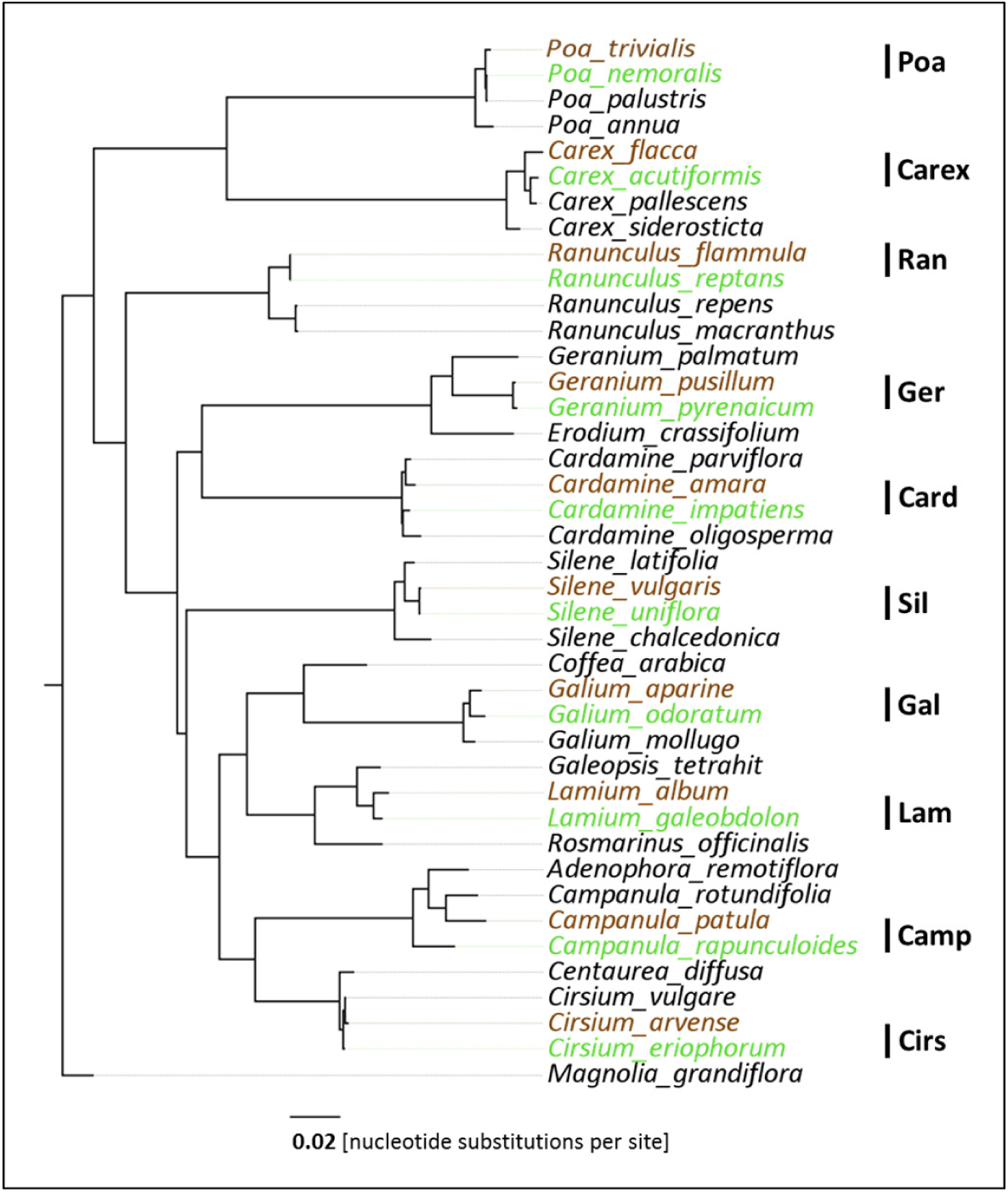
Phylogenetic relationships of the 41 species. The tree was inferred from the Bayesian analysis of the concatenated alignment of 73 plastid protein-coding genes (CDS). Posterior probability for all nodes ≥ 0.99. Brown: seed-banking (SB) species; green: nonseed-banking (NSB) species. On the right side, abbreviations for the respective SB/NSB species pairs.

## Material and Methods

### Choice of Candidate Species

Based on a large literature search (Thompson et al., 1997; Baskin & Baskin, 2014) ten quadruplets of species were chosen, each consisting in an ingroup with one seed-banking (SB) and one non seed-banking (NSB) species, and two other outgroup species (total of 40 species and one general outgroup, list in Table S1 and phylogeny in Fig. 1). Inside each quadruplet, the two ingroup species were chosen to be as closely-related and ecologically similar as possible. Details on the literature search and species classification as SB or NSB are provided in the Supplementary Text 1, and further description of species (longevity, generation time, habitat, sexual system, and pollinators) in Table S5.

### DNA Extraction, PCR and Sequencing

We sequenced the plastomes and nuclear ribosomal internal transcribed spacers (ITS) of 27 species. For 13 additional species and the outgroup *Magnolia grandiflora*, the data were downloaded from GenBank. All plant information (voucher, accession numbers) is given in Table S1. When fresh leaf material was available, we performed a chloroplast DNA (cpDNA) enrichment following a protocol (Supplementary Methods 1) adapted from (Napier & Barnes, 1995; Ostertag, 2014) to obtain sufficient yield with less plant material. For species with only herbarium specimens available or if the protoplast digestion failed (Table S1), we performed genomic DNA extraction on dried leaves using the Macherey Nagel NucleoSpin Plant II kit. All cpDNA-enriched and total genomic DNA samples were sent to GATC Biotech (Konstanz, Germany) for paired-end library construction and Illumina HiSeq sequencing.

For the polymorphism study, genomic DNA was extracted from samples of *Lamium album*, *L. galeobdolon*, *Cardamine amara*, and *C. impatiens* from different German populations. The plastid intergenic regions *rpl20*-*rps12*, *trnH*-*psbA*, and *trnL*-*trnF* + intron of *trnL* were obtained by PCR following the protocol described in (Schaefer & Renner, 2010) and Sanger sequencing by GATC (Konstanz, Germany). Sequences available in GenBank for those regions and for the plastid *trnS*-*trnG* and the nuclear ITS were added to make the sampling as geographically broad as possible. Voucher information, accession numbers and geographic origin of the plants are provided in Table S7.

### Data processing, plastid assembly and annotation

The Illumina raw data, consisting in 125-bp paired-end reads, were quality filtered using FastQC 0.11.3 (Andrews, 2010) and Trimmomatic 0.33 (Bolger *et al.*, 2014), discarding reads with average phred33 score < 20, and then *de novo* assembled with ABySS 1.9.0 (Simpson *et al.*, 2009). Plastid contigs were fished using BLASTing (blastn command, (Camacho *et al.*, 2009)) against a database of 698 plastomes available on GenBank. Further read mappings with less stringent parameters and manual inspection were necessary to extend the contigs at the borders of repeats and in low-complexity regions, and to assemble them *de novo* or based on a closely-related reference (listed in Table S1). These steps were performed using CLC Genomics Workbench 7.0.3 and Geneious R6 (Kearse *et al.*, 2012).

Eight plastomes could be assembled in one circular molecule, and ten additional were recovered at more than 98% of their length and could also be assembled because they displayed high collinearity with published plastomes of closely-related species. Finally, nine plastomes were only partially recovered, as suggested by the comparison to reference lengths and gene content (Table S1, Fig. S1). Reads were mapped on all full or partial assemblies (Table S1) with BWA (Li & Durbin, 2009) to generate two consensus for each species. The first “70%” consensus had bases replaced by IUPAC ambiguities when an alternative base was found in more than 30% of the reads. The second “70%-5x” consensus was generated from the previous one, but bases with read depth < 5 were replaced by Ns. The latter consensus was used in further phylogenetic analyses and rate calculations, whereas the 70% consensus was annotated using the Geneious annotation tool and manual verifications using the online blastn and blastp interfaces of the NCBI website (https://blast.ncbi.nlm.nih.gov/Blast.cgi) as well as the online server of tRNA-scan-SE (Lowe & Eddy, 1997). These 27 plastomes were submitted to GenBank with accession numbers provided in Table S1. Nuclear ITS sequences were obtained from all contig pools using blastn with *Corallorhiza macrantha* (GenBank accession NC_025660.1) as query.

### Phylogenetic analyses

Chloroplast protein coding genes (CDS) and chloroplast non-coding intergenic regions (NCS) were extracted from the 41 plastomes (Table S1). Single regions were aligned separately (i) by quadruplet and (ii) including all quadruplets for which the region was available + *Magnolia*, using MAFFT (Katoh & Standley, 2013) in the case of NCS, and MACSE (Ranwez *et al.*, 2011) for CDS, allowing to keep sequences in the correct reading frame. Alignments were manually inspected and discarded when they were too difficult to align (mostly in cases of non-coding regions aligned across all quadruplets) or when at least one taxon (in the case of quadruplet alignments) or 20 taxa (in the case of global alignments) were missing. For each quadruplet, all single CDS alignments were then concatenated in a total CDS matrix, and the same was done separately for NCS alignments, resulting in 10 total CDS matrices of four taxa (of 20,883 to 62,325 characters) and 10 total NCS matrices of four taxa (of 11,450 to 41,735 characters). Finally, two global CDS and NCS matrices of 41 taxa and 57,993 and 64,441 characters were obtained by concatenating the single alignments comprising all quadruplets + *Magnolia*. All single-regions and total matrices were submitted to Maximum Likelihood phylogenetic analyses based on the GTRGAMMA model implemented in RAxML (Stamatakis, 2014), with 100 bootstrap replicates. Single-region trees where compared to the trees obtained from concatenated matrices, and regions supporting with at least 70% bootstrap support a different topology than the majority where removed from the concatenated matrices for the concerned quadruplets, resulting in two new concatenated CDS and NCS matrices for each quadruplet as well as two new concatenated global CDS and NCS matrices. In addition, eleven matrices consisting of the concatenation of ITS1 and ITS2 were also generated, one for each quadruplet and a global one, and aligned with MAFFT. All concatenated matrices, including the original ones comprising conflicting regions were submitted to phylogenetic inference using MrBayes v. 3.2 (Ronquist *et al.*, 2012) under a GTR + γ model, and performing two runs of 2.5 million generations (to reach convergence). A burnin fraction of 25% was removed from the sampling before building consensus trees and analyzing posterior distributions of branch lengths (see below). The Bayesian analyses were repeated following best partitioning schemes and substitution models assessed with PartitionFinder v. 1.1.1 (Lanfear *et al.*, 2012). Finally, in order to assess the robustness of our results to an increase of sampling, CDS were also extracted from the plastomes of 358 angiosperms available in GenBank (accessions in Table S7), aligned with the CDS of our study species using MACSE, and then concatenated in a global matrix of 399 taxa and 42,132 characters. This alignment was submitted to phylogenetic Bayesian Inference using Exabayes (Aberer *et al.*, 2014) for two runs of 10 million generations with sampling frequency of 1000, which was enough to reach convergence. The most important matrices will be made available in Dryad upon acceptance of the manuscript.

### Polymorphism analyses

For the study of polymorphism in *Lamium* and *Cardamine*, each region was aligned using MAFFT and differences observed between the two sequences used in our full plastome or ITS comparisons were considered as fixed substitutions if present in all available sequences of the species, and as polymorphic if present in only some of them. Inside one quadruplet (*Lamium* or *Cardamine*), the substitutions were polarized based on the two outgroup species, and conservatively, we removed sites for which both states were found in the ingroup and which showed different states in the outgroups.

### Testing for Rate Heterogeneity and for Confounding Factors

Tajima’s tests (Tajima, 1993) for rate heterogeneity between the SB and NSB species of each quadruplet were performed on all concatenated matrices using MEGA v 7.0 (Kumar *et al.*, 2016), and taking the most closely-related species of each SB/NSB pair as outgroup, except for *Campanula*, *Cardamine* and *Ranunculus*, for which we chose *Cirsium vulgare*, *Geranium palmatum* and *R. repens*. The significance level of α = 0.05 was Bonferroni-corrected to α*_Bonf_* = 0.005 accounting for ten individual tests that were performed on each sequence data set. For each quadruplet in each Bayesian phylogenetic analysis, the relative substitution rate of SB and NSB species was obtained by estimating the post-burnin average length of their branch starting from the most recent common ancestor of both species. We also extracted the 95% confidence intervals (CI) of those branch-lengths from the post-burnin tree sample and defined a confidence interval overlap index between the SB and the NSB species to facilitate visualization of the results. This CI overlap index is defined as the difference between the lowest bound of the largest mean BL 95 % CI and the upper bound of the smallest mean BL 95 % CI normalized by the species pair mean BL. Finally, Jukes-Cantor corrected ratios of non-synonymous to synonymous substitutions (dN/dS) were calculated for each SB and NSB species in the concatenated quadruplet CDS alignments, using SNAP 2.1.1 (Korber, 2000) (www.hiv.lanl.gov).

## Results

### High quality plastome data and phylogenies

We analyse here the full plastome of 41 angiosperm species, of which 27 are generated *de novo* from species belonging to the genera *Campanula* (Campanulaceae, Asterales), *Cardamine* (Brassicaceae, Brassicales), *Carex* (Cyperaceae, Poales), *Cirsium* (Asteraceae, Asterales), *Galium* (Rubiaceae, Gentianales), *Geranium* (Geraniaceae, Geraniales), *Lamium* (Lamiaceae, Lamiales), *Ranunculus* (Ranunculaceae, Ranunculales), *Poa* (Poaceae, Poales) and *Silene* (Caryophyllaceae, Caryophyllales). We choose these genera and species because of the available detailed description of the seed-bank status and life history traits (Thompson *et al.*, 1997; Baskin & Baskin, 2014) (see details in SOM). We develop a new protocol for chloroplast DNA (cpDNA) extraction (SO Methods 1), so that DNA is extracted either 1) by enzymatic digestion of fresh leaves followed by cpDNA enrichment (yielding an average per-base read-depths of 209x), 2) directly from fresh leaves (yielding a coverage of 140x), or 3) directly from a dry herbarium specimen (yielding a coverage of 173x, Table S1). Regardless of the approach, we obtain an average per-base read depths of 199x (median of 159x) ranging from 13x (*Silene uniflora*; herbarium material) to 907x (*Galeopsis tetrahit*; cpDNA enrichment). In 19 of the 27 species, the complete plastid genome could be recovered with a high coverage (>100x, Table S1).

The quality of our 27 new plastomes and the final coverage depends on the quality of the starting material and the extraction methods. The modified enrichment protocol that we develop here works with a very limited amount of starting material, in contrast to previous work (Kolodner *et al.*, 1976; Bookjans *et al.*, 1984; Palmer, 1986; Jansen *et al.*, 2005). Compared to direct genomic DNA extraction our method yields an average increase in read depth of more than 20% and up to more than 47% when excluding species for which senescing/herbivore-damaged leaves or mainly petiole material is used (see Table S1). Intact young leaf material is best for cpDNA enrichment, but the data recovered from direct DNA extraction of 31 to 55 years-old herbarium specimens also allows us to assemble large plastome regions with high coverage.

We find that intra-plastome read depth is heterogeneous across our samples but of sufficient quality for read mapping and substitution calling (Table S1). Out of 27 plastomes we find eight with a sufficient coverage to be completely assembled. Ten other species require a few junctions between non-overlapping contigs via inference of gaps (up to ten gaps) using high collinearity to available references. The cumulated length of these represents between 98% and 100% of the reference length suggesting that those plastomes are in fact (almost) complete. Finally, low coverage often combined with the presence of repeated sequences prevent the complete assembly of nine plastomes (*Galium odoratum*, *Campanula sp.*, *Carex sp.*, and two *Geranium* species, Table S1). For those nine species, we still recover cumulated lengths between 51% and 83% of their respective reference lengths with most of the missing nucleotides being non-coding DNA.

The plastomes of all taxa studied here have a conserved gene content with 78 protein-coding genes, four rDNAs and 30 tRNAs present in most lineages (Fig. S1). Some missing genes have a distribution consistent for all species of a genus, suggesting that their absence is not due to gaps in our sequencing but to a loss of function (details in methods). The quadripartite plastome structure consisting in two large (LSC) and small (SSC) single copy regions separated by one inverted repeat (IR), as well as gene collinearity are conserved in most lineages, and thus support the homology of most inter-genic regions used in the following phylogenetic analyses. The consistency of results is checked 1) between coding sequences (CDS), non-coding sequences (NCS) and the nuclear ITS locus, 2) between partitioned datasets with different substitution models across genes and non-partitioned datasets, and 3) between datasets where genes producing trees conflicting with the most often recovered topology were removed and datasets where all genes were kept (see methods).

We build a phylogeny of ten angiosperm clades spanning nine orders (listed above) by including quadruplets of species consisting of one with well documented long term persistent seed-bank (hereafter SB), one with a relative absence of seed-bank compared to its sister (NSB), and two outgroups belonging to the same genus or a closely-related one. After controlling for supported incongruences between gene trees (see methods), three times ten local alignments of four taxa and three global alignments of the ten clades are built concatenating separately protein-coding plastid genes (CDS), non-coding plastid (NCS) regions, and nuclear ITS data. We then compute the phylogenetic trees using Bayesian methods and find that all concatenated data sets yield qualitatively identical, well-supported (most Bayesian posterior probabilities PP = 1), tree topologies (Fig. 1), regardless of species sampling, partitioning, or presence or absence of conflicting regions (Fig. S2, details in methods). The relationships between the families are consistent with those presented by the Angiosperm Phylogeny Group (APGIV, 2016) except for a conflict between the positions of *Silene* (Caryophyllaceae) as sister to Geraniaceae+Brassicaceae in the non-coding (NCS) trees (Fig. S2c,d), and sister to Asteraceae+Lamiaceae+Rubiaceae in the CDS trees (Fig. 1; Fig. S2a,b), the latter being the position accepted by APGIV (2016). Note that the tree based on the nuclear ITS dataset is less well resolved, with many low-supported (PP < 0.97) clades (Fig. S2e and details of conflicts in SOM). Finally, as intra-generic topologies of *Campanula*, *Cardamine* and *Ranunculus* appear different from initially expected, we redefine outgroups for those genera (see methods).

### Seed-banking species have equal or higher substitution rates than non-seedbanking species

Based on these phylogenies we obtain the branch length (BL) measures BL_SB_ and BL_NSB_ respectively for seed-banking and non-seed-banking species since they diverged from their most-recent common ancestor. Our principal result is that the substitution rate in seed-banking (SB) species is equal or higher than that of non-seed-banking (NSB) species, namely BL_SB_ > BL_NSB_ This result is supported in analyses based on partitioned or non-partitioned alignments including all regions or only non-conflicting ones (conflicting regions for the phylogeny building), and including all ten clades or only local alignments for quadruplets. All branch lengths for SB and NSB species estimated in the different analyses are reported in Tables S2 and S3.

In Fig. 2, we show the result of the ratio BL_SB_ / BL_NSB_ for non-partitioned analyses at the angiosperm level. If seed-banking has no effect on the rate of substitution the ratio BL_SB_ / BL_NSB_ is expected to be one, while most genera show a higher rate of evolution of the seed-banking species (Fig. 2). For both plastid CDS and NCS, eight out of ten species pairs display longer branch lengths in the SB than in the NSB species (sign test p-value = 0.044), and results are consistent per quadruplet for CDS and NCS. Two exceptions to this consistency are seen with *Galium* and *Ranunculus*, while *Geranium* is the only quadruplet showing BL_SB_<BL_NSB_ both at CDS and NCS (Fig. 2a,b). Note that in all these cases, the 95 % confidence interval (CI) overlap index is always positive, indicating significant differences in substitution rate in all pairs for both CDS and NCS datasets (black rectangles in Fig. 2a,b). Additionally, we perform Bonferroni-corrected Tajima’s tests for rate heterogeneity (p-values indicated in Table 1), which shows that four (*Campanula*, *Cirsium*, *Lamium*, and *Poa*) out of ten species SB/NSB pairs have significantly different substitution rates (p-value < 0.005), in a consistent manner between CDS and NCS datasets. The inconsistency observed for *Galium* and *Ranunculus* between CDS and NCS in Fig. 2 is resolved in favour of the result at NCS because the p-values from the Tajima’s tests and the CI overlap index are respectively lower and higher for NCS than CDS for both quadruplets. We favour thus BL_SB_>BL_NSB_ in *Galium* and BL_SB_<BL_NSB_ in *Ranunculus* (Fig. 2, Table 1). Nuclear ITS data confirm for most quadruplets the higher rate of substitution in seed-banking versus nonseed-banking species (Fig. 2c) although topological conflicts prevent the use of *Cardamine* and lead us to use genus-level data for *Poa* (Fig. S2e-g). The only conflicts observed between plastid and ITS results are for *Carex* and *Ranunculus*, but these are not supported by CI overlap indexes, or by Tajima’s test p-values. We thus conclude that both genera follow their plastid global trend, i.e. BL_SB_>BL_NSB_ (Fig. 2 and Table 1). Finally, we confirm our results from Fig. 2 using a phylogenetic tree for only CDS with 399 available taxa from most angiosperm families and orders (Fig. S2h). The results are confirmed for most quadruplets, including *Cirsium*, *Geranium* and *Cardamine* for which a closer outgroup was available in this extended dataset. The only exception is *Silene*, in which the non-seed-banking *S. uniflora* has a slightly longer branch than the seed-banking *S. vulgaris* (Fig. S2h), a result contradicting those from Fig. 2a (Table S3, Fig. S2a-e).

**Figure 2.**
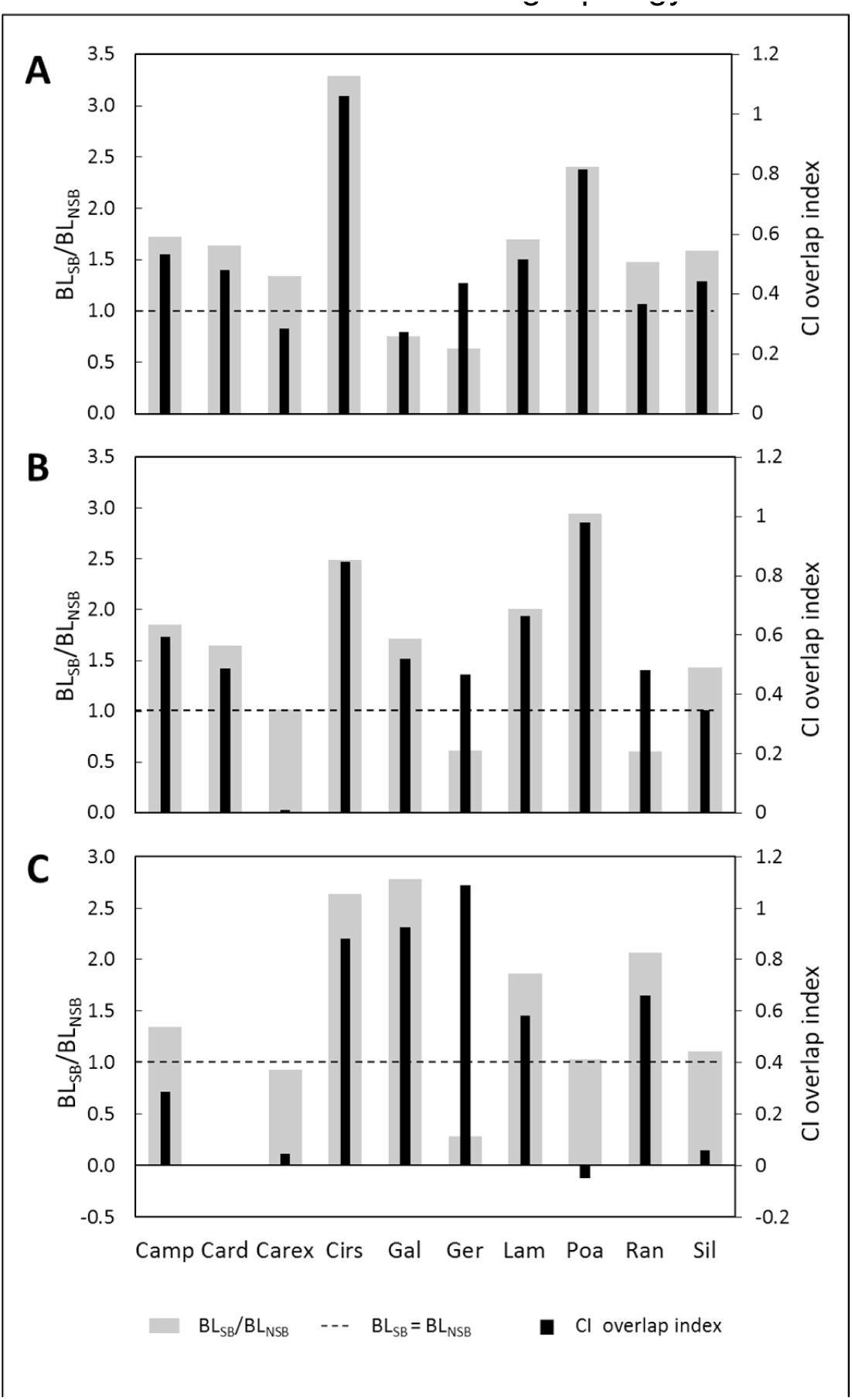
Relative substitution rates of SB/NSB based on branch lengths (BL). (a). Chloroplast coding sequences (CDS), (b). chloroplast non-coding sequences (NCS) and (c). nuclear rDNA ITS sequences. The grey bars indicate the average BL ratios over all post-burnin sampled trees (scaled by primary y-axis). The dashed lines indicate the ratio BL_SB_/BL_NSB_=1. The second statistics shown is the 95 % confidence interval (CI) overlap index, plotted as black bars and with the second y-axis. A positive CI overlap index indicates significantly different substitution rates. Data for Cardamine nuclear ITS were not included into the statistic evaluation due to conflicting topology.

**Table 1.** P-values of the Tajima’s test for rate heterogeneity, in bold when BL_SB_ significantly superior to BL_NSB_ after Bonferroni correction. The test was performed on concatenated sequence alignments. For *Ranunculus* the outgroup species used for ITS (*R. venetus*) differed from the one used for CDS and NCS (*R. repens*).

|  | CDS | NCS | ITS |
| --- | --- | --- | --- |
| *Campanula* | **0.000** | **0.000** | 0.104 |
| *Cardamine* | 0.396 | 0.243 | 0.011 |
| *Carex* | 0.006 | 0.393 | 0.465 |
| *Cirsium* | **0.000** | **0.000** | 0.028 |
| *Galium* | 0.029 | 0.005 | **0.001** |
| *Geranium* | 0.190 | 0.010 | 0.257 |
| *Lamium* | **0.000** | **0.000** | 0.695 |
| *Poa* | **0.000** | **0.000** | 0.414 |
| *Ranunculus* | 0.317 | 0.128 | 0.317 |
| *Silene* | 0.174 | 0.251 | 1.000 |

### Life-history traits or selection do not explain the difference in substitution rates

We control here that the differences we observe between BL_SB_ and BL_NSB_ are not due to confounding life-history traits or differential selection acting on chloroplast genomes. We document for each species pair the estimated relative generation time, location, habitat disturbance or sexual system as potential explanatory factor (Table S5). Location, habitat disturbance or sexual system (Table S5) are widely similar between sister species and thus unlikely to explain the higher substitution rates in SB. There is no significant correlation between BL estimates (or their ratios) and generation time (Fig. 3a, Table S4). Generation time is here fairly heterogeneous and difficult to estimate as most species are perennial. The absence of correlation is chiefly due to the fact that in some cases the SB species has a higher generation time than the NSB, while the reverse is true in other cases. The substitution rate difference cannot be attributed either to the single-species average plastome coverage (Fig. S3a,b). An alternative explanation could be the difference in selective pressure (positive or purifying) acting on chloroplast genomes of seed-banking or non-seed-banking species. However, dN/dS values do not correlate with BL (Fig. 3b), or with relative generation times (Fig. S3c). Interestingly, the dN/dS ratios appear similar between sister species but different across genera (Table S6), indicating that the strength of positive or purifying selection is defined rather by genus wide constraints (such as genomic context, habitat or local selective pressure) than at the species specific level or due to SB or NSB traits.

**Figure 3.**
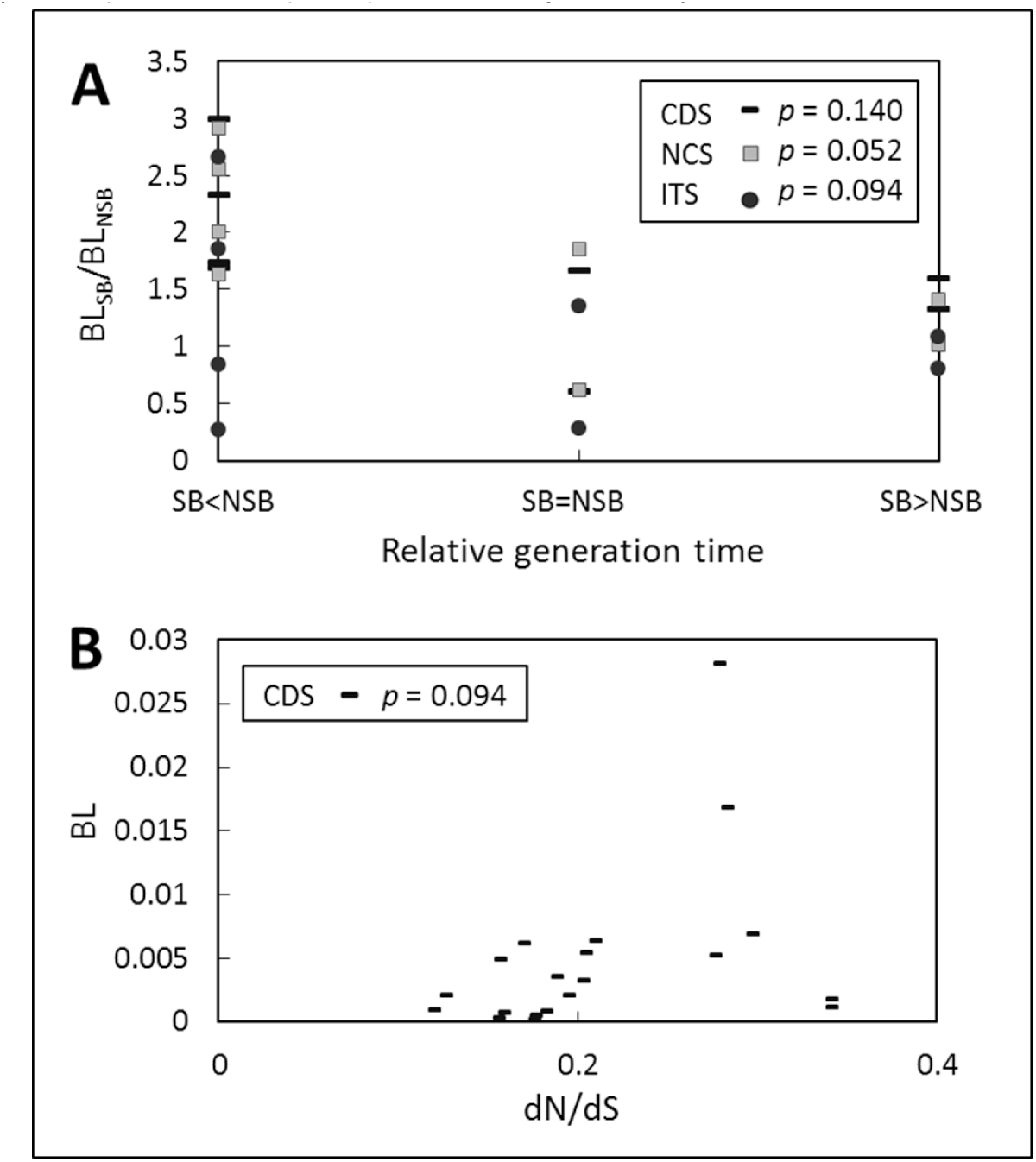
Correlation between substitution rates and additional factors. (a). BL_SB_/BL_NSB_ ratios as a function of relative generation times of SB and NSB species. (b). Branch length per species (BL) as a function of dN/dS ratio. Data were obtained from non-partitioned concatenated sequence alignments. P-values given in the inset boxes result from two-tailed t-tests for significant pairwise correlation (see table S4) and relate to CDS (bars), NCS (squares) and ITS (dots) data, respectively.

### Polymorphism does not explain the differences in substitution rates

A final possible explanation for the difference in substitution rate between seed-banking and non-seed-banking species lies in the amount of polymorphism. As SB species may exhibit higher rates of polymorphism than NSB, our substitution rate could be affected by such effect. We thus investigate the relationship between polymorphic sites and substitutions in four plastid intergenic spacers, one plastid intron and the nuclear ITS of SB and NSB species of *Lamium* and *Cardamine* (Table 2). Additional sequences from different locations in Germany and abroad (Table S7) reveal that in both lineages the higher number of substitutions in the seed-banking-species (reported in Fig. 2 and Table S3) consists indeed mostly of fixed mutations. Following the theory of storage effect, seed-banking species exhibit nevertheless a higher number of polymorphic sites than non-seed-banking species. In the five plastid regions grouped together, 33% of observed substitutions in the seed-banking *L. album* are in fact polymorphisms against 8% in the non-seed-banking *L. galeobdolon* and a similar outcome is found for *Cardamine* (5% against 0%). This is also the case in the nuclear ITS for *Cardamine* (33% against 25%) but not in *Lamium* (0% against 22%).

**Table 2.** Number of substitutions and polymorphisms in plastid and nuclear regions of *Lamium* and *Cardamine*. Counts of the number of nucleotides as substitutions (F, fixed) or as polymorphism (P) in the two seed-banking (SB) and non-seed-banking (NSB) species of the given genus.

|  | Length in bp<br>*Lamium* /<br>*Cardamine* | *L.*<br>*album*<br>(SB) |  | *L.*<br>*galeobdolon*<br>(NSB) |  | *C. amara*<br>(SB) |  | *C.*<br>*impatiens*<br>(NSB) |  |
| --- | --- | --- | --- | --- | --- | --- | --- | --- | --- |
|  |  | F | P | F | P | F | P | F | P |
| **Plastome** | 2068 / 1638 | 16 | 8 | 12 | 1 | 20 | 1 | 6 | 0 |
| **Nuclear**<br>**ITS** | 448 / 453 | 16 | 0 | 7 | 2 | 2 | 1 | 6 | 2 |

## Discussion

### Seed-bankers have equal or higher substitution rates

Our main result shows that substitution rates in seed-banking species are equal or higher than in species with less persistent seed-banks, as observed for chloroplast genome protein-coding and non-coding sequences and consistently across various groups of angiosperms. The results for the nuclear ITS region are less clear but seem to show a similar trend, which is also supported by the preliminary work of Whittle (Whittle, 2006) on other angiosperm genera. The broad choice of studied genera is possible because we also develop a protocol for chloroplast DNA enrichment for high throughput DNA sequencing. This allows to analyze a unique combination of plastomes from yet unstudied seed-banking and non-seed-banking species from small amount of fresh starting material, including senescent of herbivore-damaged leaves.

The detrimental effect that prolonged seed soil storage may have on DNA integrity (Levin, 1990; Waterworth *et al.*, 2016) is expected to result in generally elevated mutation rates in SB species compared to closely related NSB species. We use here predictions about the molecular clock hypothesis (Zuckerkandl & Pauling, 1965), stating that the neutral substitution rate is a direct function of the nucleotide mutation rate and does not depend on the species size (Kimura, 1968). Since life histories of our study species are largely identical within SB/NSB pairs (Table S5), the equal or even higher number of substitutions in the SB species stems from the seed-bank. We here therefore reject the alternative hypothesis that mutations do not occur in seeds. In addition, we conclude that the absence of noticeable differences in selection within our species pairs (Table S6) indicates a similar proportion of effectively neutral (or nearly neutral) mutations (Ohta & Kimura, 1971) in both SB and NSB species. In other words, the proportion and selective coefficients of deleterious mutations arising in the seeds appear to be similar to those of the mutations originating in above-ground plants. The seed-bank acts therefore as source of additional molecular variants whose likelihood of getting fixed in the population is similar to those arising in above-ground plants.

A meta-analysis of 13 studies (Honnay *et al.*, 2008) and studies of genetic diversity in *A. thaliana* populations in Norway (Lundemo *et al.*, 2009; Falahati-Anbaran *et al.*, 2014) do not show, however, a significant accumulation of rare genotypes in the soil seed-bank. This apparently contradicts the notion of persistent seed-banks as sources of genetic variability. We explain these results by a moderate proportion? of new fitness-neutral genotypes which constantly germinate from the seed-bank and homogenize above- and below-ground populations genetically. This is consistent with the fact that disrupted genome integrity with respect to DNA double strand breaks does impair germination in *A. thaliana*, while the effects of moderate chemical mutagenesis are largely tolerated (Waterworth *et al.*, 2010). This suggests first that the majority of persisting seed-bank-borne mutations would thus be neutral or nearly neutral (Ohta & Kimura, 1971) and the selection against mutant seeds in early life stages (seedling) may be less severe than previously assumed. Second, environmental uncertainties against which seed-banks constitute a bet hedging strategy (Brown & Venable, 1986; Evans & Dennehy, 2005) may result during certain periods in stronger random genetic drift due to catastrophic decrease in above-ground population size. This might allow for establishment and propagation of seedlings emerging from mutant seeds that would likely be outcompeted under normal environmental conditions, allowing mutant alleles to rise in frequency. Our results thus call for studies on the natural variation and fitness effect of seed born mutations.

### Seed-banking influences micro- and macro-evolutionary studies

Assuming a neutral model of evolution, the amount of genetic polymorphism and ecological data on the average above-ground population size can be jointly used to infer the persistence of the seed-bank (Tellier *et al.*, 2011) and past demographic events (Živković & Tellier, 2012). Ignoring the mutation rate in seeds thus yields errors in the estimations of the length of coalescent trees, and as a consequence on the estimation of neutral population parameters such as the effective population size and past demographic history. In addition, with regards to natural selection, seed-banks are also a source of potentially advantageous mutations which increase the adaptive potential of the population, though note that a persistent seed-bank has for effect to slow down the speed of natural selection (Hairston Jr & De Stasio Jr, 1988; Koopmann *et al.*, 2016) compared to populations with short seed-banks. Finally, when studying the recent divergence between population and/or species, taking into account the adequate rate of mutation (including in seeds) is important to estimate times of splits, as seed-banks tend to decrease differentiation (Vitalis *et al.*, 2004) and increase incomplete lineage sorting. An example of incomplete lineage sorting caused by recent divergence and possible persistent seed-banks can be seen in wild tomato species (*Solanum* clade, (Pease *et al.*, 2016)).

In phylogenetics substitution rate heterogeneity is well-known to occur not only between and within genes but also between taxa at a given gene, from inter-family to inter-species taxonomic levels (Thomas *et al.*, 2006). The factors generating this heterogeneity can be differences in 1) metabolic rates, 2) life-history such as generation time, and 3) altitude and latitude (Bromham *et al.*, 2015). As it is especially difficult to disentangle the effects of these various factors on substitution rate heterogeneity between lineages, phylogeneticists use models of nucleotide evolution allowing substitution rates to change during the process of cladogenesis and lineage evolution. Most of the models assume rate autocorrelation, that is assume *a priori* that large jumps of the rate along the phylogeny are less probable than gradual changes (reviewed in (Ho & Duchêne, 2014)). Genetically-dependent rate heterogeneity is commonly expected to be more autocorrelated than changes in environment-dependent rates. We here suggest that *a priori* knowledge of lineages with or without long-term seed-banking is extremely useful for phylogeny estimation, for example to further refine models dealing with the molecular dating of angiosperm phylogenies. As a corollary, seed-banking persistency could be predicted for populations exhibiting different rates if other substitution rate-modifying factors can be ruled out.

A major limitation in studying seed-banking is the availability of reliable data on this trait, especially regarding the absence of seed-banking. Seed-bank persistence is in effect a difficult to measure and gradually expressed trait, and we resort here to comparing closely related species with a pronounced difference in seed-bank longevity. Without the many small-scale ecological studies performed during the last century our analysis would have not been possible (compiled in (Thompson *et al.*, 1997; Baskin & Baskin, 2014)). Seed-banks are of importance for conservation biology and restauration ecology, and more data on seed longevity and seed-banking in angiosperms are also needed to better understand macro- and micro-evolutionary processes. Our study shows that the seed reservoir is not genetically inert and exhibit new mutations measured here as substitution rates, which should be taken into account in ecological as well as population genetic and phylogenetic analyses.

## Acknowledgements

We thank A. Saatkamp (Aix Marseille University - IMBE, France) for his initial advises and for helping us to access the Thompson database, and G. Achaz for comments on the manuscript. AT is supported by the Deutsche Forschungsgemeinschaft grants TE809/1-1, TE809/7-1 and sequencing was funded in part by the Federal Ministry of Education and Research (BMBF, Germany) within the AgroClustEr Synbreed - Synergistic plant and animal breeding (grant 0315528I).

### Author Contributions

MD, SB, HS and AT designed the study and wrote the manuscript. MD and SB performed the analyses, MD and HS collected samples, MD, SS and SB performed the lab work.

## Supporting Information

Mutation rates in seeds and seed-banking influence substitution rates across the angiosperm phylogeny

*Marcel Dann, Sidonie Bellot, Sylwia Schepella, Hanno Schaefer, Aurélien Tellier*

The following Supporting Information is available for this article:

**Methods S1:** Chloroplast (cp) DNA enrichment protocol.

**Text S1.** Evidence SB/NSB annotations and expected candidate species clade topologies.

**Table S4.** Summary of correlation analyses between BL and additional factors.

**Table S6.** Estimate of natural selection acting on protein-coding chloroplast genes.

**Figures S1, S2.** attached as pdf

**Table S3.** Correlation analyses of substitution rates as a function of additional factors

**Tables S1-S3, S5, S7.** attached as excel files.

## METHODS S1: Chloroplast (cp) DNA enrichment protocol

From 0.26 to 3.10 g fresh leaf material was harvested from living plants preferring the youngest leaves available. Leaves were cleaned with de-ionized water thoroughly and cut into pieces with a razor blade inside a 9 cm petri dish. Leaf fragments were covered with 15 ml enzyme solution immediately (see Tables below for the composition of all solutions and description of the equipment), making sure all fragments were coated with liquid completely. Petri dishes were sealed with laboratory film and placed on a rotary plate (Orbit LS, Labnet) at 60 rpm at room temperature (~22 °C) for 4 hours.

In principle, chloroplast isolation from protoplasts was achieved as described by Napier and Barnes (1995). Protoplasts were collected by passing the leaf + enzyme solution through a nylon net (pore size 140 μm) and flushing the petri dish with 10 ml cold WIMK. The filter was flushed with another 10 ml cold WIMK to minimize protoplast loss.

The protoplast solution was collected in 50 ml falcon tubes and centrifuged at 4 °C and 3988 rcf for 5 min in a SIGMA 2-16K refrigerated centrifuge (SIGMA Laborzentrifugen GmbH). The supernatant was discarded and protoplasts were re-suspended in 10 ml cold WIMK. The suspension was centrifuged at 4 °C and 3988 rcf for 5 min in a swing-out rotor and the supernatant discarded. Protoplasts were re-suspended in 1.5 ml cold GRM and transferred into 2 ml reaction tubes.

To fragment intact protoplasts, the solution was drawn up in a 5 ml syringe through a 0.45 mm cannula and re-ejected eight to twelve times. The solution of fragmented protoplasts was put onto an ice-cooled 40 % / 80 % Percoll^®^-in-GRM step gradient. Percoll^®^ gradients were prepared in 15 ml falcon tubes by carefully placing 2 ml of 80 % Percoll^®^ suspension under 4.5 ml of 40 % Percoll^®^ suspension with a Pasteur pipette. After centrifugation for 20 min at 4 °C and 3988 rcf in a swing-out rotor the supernatant was removed using a 1 ml pipette. Intact chloroplasts were collected from the 40-80 % interface with a Pasteur pipette avoiding uptake of the 80 % phase. The chloroplast fraction was split into aliquots of a maximum 750 μl and transferred into 2 ml test tubes, which were then filled up to 2 ml with cold GRM. The solution was mixed thoroughly and centrifuged at 2000 rcf for 5 min in a benchtop centrifuge (Centrifuge 5424, Eppendorf AG). The supernatant was removed and the pellet was re-suspended in 0.5 ml cold GRM and chloroplasts were pelleted at 2000 rcf for 5 min in a benchtop centrifuge.

Finally, cpDNA extraction was performed with the Macherey Nagel NucleoSpin Plant II kit following the manufacturer’s suggested protocol, using the chloroplast pellet as starting material.

**Solutions, chemicals and enzymes used for chloroplast enrichment:**

| Solution | Ingredients | Final Concentration | Product ID | Company |
| --- | --- | --- | --- | --- |
| **Enzyme Solution** | D-Mannitol | 400 mM | 4175.1 | Carl Roth |
|  | CaCl<sub>2</sub> anhydr. | 8 mM | CN93.3 | Carl Roth |
|  | MES | 5 mM | 4256.2 | Carl Roth |
|  | Cellulase | 1 % w/v | C1184 | Sigma-Aldrich |
|  | Pectinase | 0.25 % w/v | 17389 | Sigma-Aldrich |
|  | BSA | 0.1 % w/v | 0052.1 | Carl Roth |
| **WIMK** | D-Mannitol | 500 mM | 4175.1 | Carl Roth |
|  | MES | 5 mM | 4256.2 | Carl Roth |
| **GRM**<br>**(Grinding Medium)** | D-Sorbitol | 330 mM | 6213.1 | Carl Roth |
|  | HEPES | 50 mM | HN78.1 | Carl Roth |
|  | EDTA | 2 mM | CN06.3 | Carl Roth |
|  | MgCl<sub>2</sub> anhydr. | 2 mM | KK36.1 | Carl Roth |
|  | MnCl<sub>2</sub>*(H<sub>2</sub>O)<sub>4</sub> | 1 mM | T881.3 | Carl Roth |

## TEXT S1: Evidence SB/NSB annotations and expected candidate species clade topologies

### Expected clade topologies

Extended clade tree topologies (original SB/NSB/outgroup 1 plus outgroup 2 obtained from GenBank) according to published phylogenies.

### Comparative seed bank persistence

Candidate species seed banking properties were compared based on two literature-derived parameters: 1) the seed longevity index, which is defined as ratio of N[STP+LTP]/N[transient+STP+LTP] seed banking records (1), and 2) the proportion of LTP seed banking records (i.e. N[LTP]/N[transient+STP+LTP] records). Both indices were calculated based on the Thompson data base entries (2) and their respective numerical values are listed in table S3. Seed banking properties of species (-pairs) for which these parameters proved ambivalent and did not allow for clear SB/NSB assignments are presented in more detail below.

## Campanula

**Expected topology:** (((*C. patula*, *C. rapunculus*) *C. rotundifolia*) *Adenophora spec.*) (3)

### Campanula rapunculoides

Few records exist for *C. rapunculoides* seed banking traits. Thompson et al. (1997) gathered two records implying a transient or short term persistent seed bank, respectively. A more recent study confirmed short term seed bank persistence with documented buried seed longevity for ≤3 years (4). Moreover the existence of tuberoid storage organs implies reliance onto another form of bet-hedging against spatio-temporal habitat variability, which was found to correlate negatively with soil seed bank persistence ((5) and references within). Thus *C. rapunculoides* can be assumed to be a relatively weak seed banker when compared to *C. patula* (33 % LTP; 50 % persistent records) and *C. rotundifolia* (4 % LTP; 22 % persistent records; see (2)).

## Cardamine

Expected topology 1: (((*C. amara, C. oligosperma*) *C. parviflora*) *C. impatiens*) (6)

Expected topology 2: (((*C. amara, C. parviflora*) *C. oligosperma*) *C. impatiens*) (6)

### Cardamine impatiens

To us no recent records of *C. impatiens* seed bank persistence are known. Thompson et al. (2) list one transient and one short term persistent record, respectively. Additional indication of *C. impatiens* being a transient/short term seed banker are provided by observations of high seed density in the soil (7, 8) while population density was observed to be considerably fluctuant among years (9).

## Carex

**Expected topology:** ((*C. acutiformis*, *C. pallescens*) *C. flacca*) *C. siderosticta* ? (10, 11)

### Carex acutiformis

While most sedges are considered (long term) persistent seed bankers, ecological data available makes *C. acutiformis* appear as an exception. Several records confirm frequent occurrence in aboveground vegetation while soil seed bank density is low (12) or empty (13, 14). In general seed bank density was observed to be relatively low when compared to other sedges (reviewed in (12)). In fact, to our knowledge, there is but one record of *C. acutiformis* seeds staying viable for >15-20 years; the seven remaining records summarised by Thompson et al. (2) imply transient to short term persistent seed banking. The original paper (…) documenting such long seed longevity is not available to us, however. More recent records postdating the Thompson et al. database release confirm the short term persistence of *C. acutiformis* seed banks (15) and imply reliance on clonal growth rather than sexual reproduction (12, 16). In more recent studies seed output was shown to be low and germination from seeds could not be observed at all (17). Schütz (15) observed low viable seed bank formation (>90 % of buried seeds dead or fatally germinated) under conditions favourable for germination and near-complete germination of surface-sown seeds within 1 year, while germination rates were very low under non-favourable conditions (15, 18). In field studies the germination rate was found to be rather low in general as well (16). While Schütz concluded from that that “the ability to form long-persistent seed banks is obvious”, rapid loss of *C. acutiformis* seed viability (19) and the overall low seed output make the formation of an effective, long term persistent seed bank appear unlikely. Moreover in recent publications it has become common practice to categorize *C. acutiformis* as transient (20) or short term persistent seed banker (21-23).

### Carex flacca

*Carex flacca* has been found to produce a high number of seeds resulting in high seed bank densities (4, 24) and could be shown to form a LTP seed bank repeatedly (> 20 yrs listed by Thompson et al.(2), > 39 yrs (25)) and is accepted as a LTP seed banker (26).

## Cirsium

Expected topology*: ((*C. eriophorum*, *C. vulgare*) *C. arvense*) *Centaurea spec.*) (27) *\* derived from morphological and life history data!*

## Galium

**Expected topology:** (((*G. aparine*, *G. odoratum*) *G. mollugo*) *Coffea spec.*) (28)

### Galium aparine

*Galium aparine* does not tend to form a long term persistent seed bank *sensu* Thompson on a regular basis, but short term soil seed bank persistence up to 5 years has frequently been reported (2). Considerable seed incorporation into deeper soil layers and a resulting delay in seed germination for several seasons has been observed for this species (29-31). In addition maximum seed longevity of 7-8 years for soil surface storage has been reported (32) (cited in (30)). Hence *Galium aparine* can legitimately be labelled as seed banking species when compared to *Galium odoratum*.

### Galium odoratum

*Galium odoratum* has consistently been found to be a transient seed banker *sensu* Thompson (2, 33). Seed germination itself could rarely be observed (34) and despite being frequent in the aboveground vegetation absence from the soil seed bank has been reported repeatedly (29, 35). Reproduction was found to be mainly vegetative and seed production dispersal to be rather poor (33, 36). Taken together the available evidence leaves little doubt about *Galium odoratum* forming a strictly transient seed bank, if any.

## Geranium

**Expected topology:** (((*G. pusillum*, *G. pyrenaicum*) *G. palmatum*) *Erodium spec.*) (37)

### Geranium pyrenaicum

Few records on seed bank persistence are available for *Geranium pyrenaicum*. Transience (3-12 months (38)) and short term persistence (≤4 years (2)) have been reported once, respectively. However, additional ecological data suggests *Geranium pyrenaicum* not to be a long term seed banking species. First of all low seed density in soil has been reported (39). Germination requirements have been found to be very unspecific (40), while average germination rate has been found to be close to unity (94 % (41)). Finally *G. pyrenaicum* has been found to rather match the profile of a *K*-strategist, while *G. pusillum* represents an *r*-strategist (42). This annotation implies *G. pusillum* to form a more pronounced and persistent seed bank when compared to *G. pyrenaicum* (43).

## Lamium

**Expected topology:** (((*L. album*, *L. galeobdolon*) *Galeopsis spec.*) *Rosmarinus spec.*) (44)

## Poa

**Expected topology:** (((*P. nemoralis*, *P. palustris*) *P. trivialis*) *P. annua*) (45)

## Ranunculus

**Expected topology:** ((*R. flammula*, *R. reptans*) *R. repens*) *R. macranthus* ? (46, 47)

### Ranunculus reptans

Seed bank persistence records for *R. reptans* are scarce, corresponding to the species rarity. One record listed by Thompson et al. (2) suggests short term persistence. A more recent observation implies *R. reptans* not to from a soil seed bank that buffers the population from environmental hazards (48). *R. reptans* seems to rely primarily on clonal reproduction (49, 50), which is suggested to be owed to self-incompatibility and vegetation periods too short for successful fruiting and seed set (51, 52). Hence we labelled *R. reptans* as a non-seed banking species in this study, contrasted by *R. flammula* for which long term persistent seed banking records are abundant (2).

## Silene

**Expected topology:** ((*S. uniflora*, *S. vulgaris*) *S. latifolia*) *S. chalcedonica* ? (53)

### Silene uniflora

To our knowledge no explicit seed bank persistence records are available for *Silene uniflora*. In one study conducted *S. uniflora*, while present aboveground, was found absent from the soil seed bank (54). In addition ecological data suggests *S. uniflora* to be a non-seed banking species. Average seed production was reported to be very low while seed predation is prevalent (55), suggesting little capacity to form large seed banks. Moreover seed germination time polymorphism was reported among but not within maternal families (56, 57), implying *S. uniflora* doesn’t rely on a soil seed bank for bet hedging against environmental uncertainties (58). Finally populations reportedly grow on very shallow and poor substrate (59), indicating particularly bad abiotic conditions for soil seed bank formation.

---

**Table S4.**
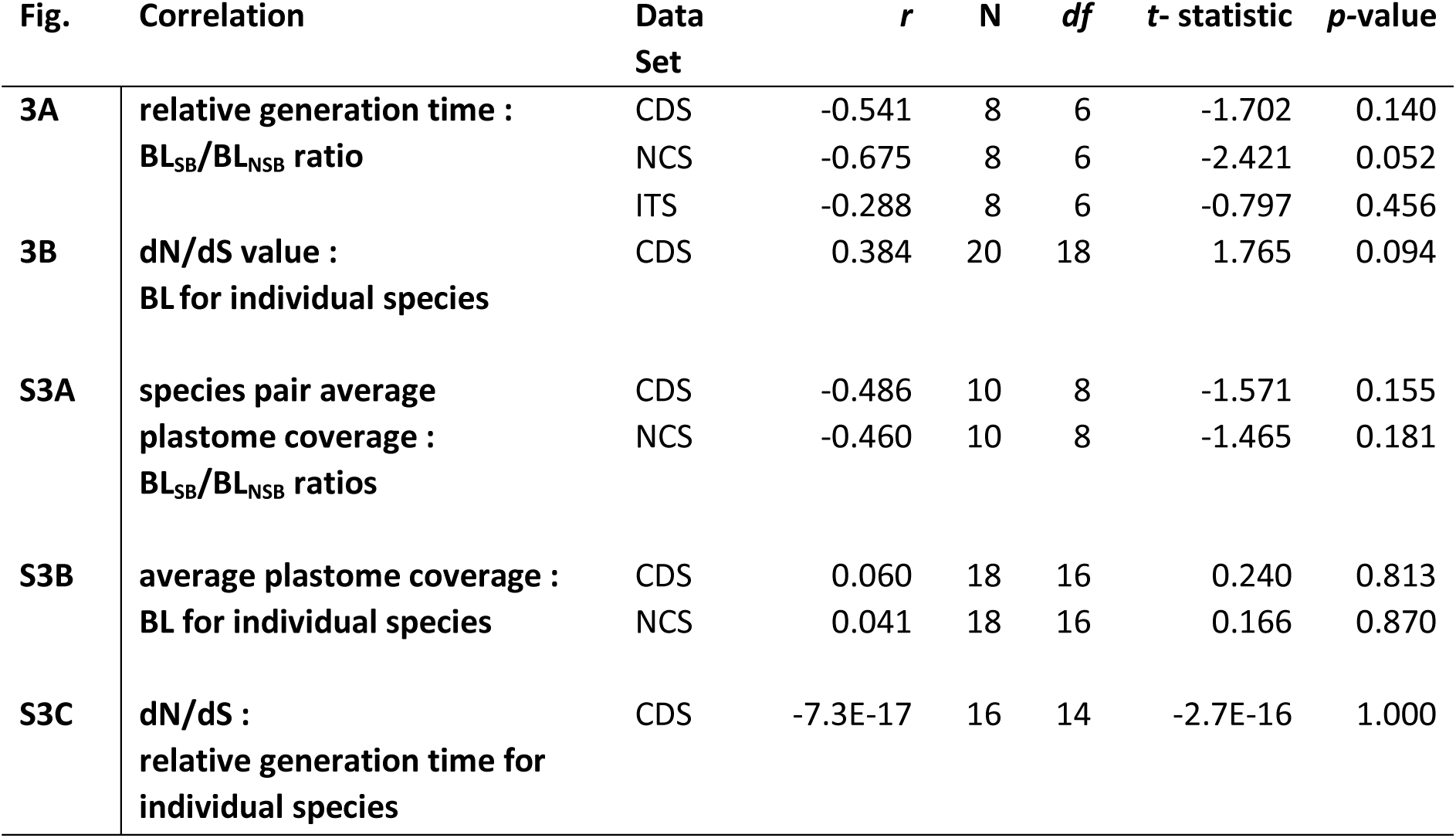
Summary of correlation analyses between BL and additional factors. The correlations coefficients from Figure S3 are given here. Pearson’s correlation coefficients, sample sizes and degrees of freedom are indicated as r, N and df, respectively. T-statistics and p-values correspond to two-sided t-tests for significant correlation.

| **Fig.** | **Correlation** | **Data Set** | ***r*** | ***N*** | ***df*** | ***t*- statistic** | ***p*-value** |
| --- | --- | --- | --- | --- | --- | --- | --- |
| **3A** | **relative generation time :** | CDS | -0.541 | 8 | 6 | -1.702 | 0.140 |
|  | **BL<sub>SB</sub>/BL<sub>NSB</sub> ratio** | NCS | -0.675 | 8 | 6 | -2.421 | 0.052 |
|  |  | ITS | -0.288 | 8 | 6 | -0.797 | 0.456 |
| **3B** | **dN/dS value :** | CDS | 0.384 | 20 | 18 | 1.765 | 0.094 |
|  | **BL for individual species** |  |  |  |  |  |  |
| **S3A** | **species pair average** | CDS | -0.486 | 10 | 8 | -1.571 | 0.155 |
|  | **plastome coverage :** | NCS | -0.460 | 10 | 8 | -1.465 | 0.181 |
|  | **BL<sub>SB</sub>/BL<sub>NSB</sub> ratios** |  |  |  |  |  |  |
| **S3B** | **average plastome coverage :** | CDS | 0.060 | 18 | 16 | 0.240 | 0.813 |
|  | **BL for individual species** | NCS | 0.041 | 18 | 16 | 0.166 | 0.870 |
| **S3C** | **dN/dS :** | CDS | -7.3E-17 | 16 | 14 | -2.7E-16 | 1.000 |
|  | **relative generation time for individual species** |  |  |  |  |  |  |

**Table S6.** Estimate of natural selection acting on protein-coding chloroplast genes. Ratios of non-synonymous to synonymous substitutions (dN/dS) from CDS alignments. Subscript SB and NSB indicate seed-banking and non-seed-banking species, respectively.

| | dN/dS | $\frac{(dN/dS_{SB})}{(dN/dS_{NSB})}$ |
| --- | --- | --- |
| *Campanula patula*<sub>SB</sub> | 0.279 | 0.984 |
| *Campanula rapunculoides*<sub>NSB</sub> | 0.284 |  |
| *Cardamine amara*<sub>SB</sub> | 0.205 | 1.006 |
| *Cardamine impatiens*<sub>NSB</sub> | 0.204 |  |
| *Carex flacca*<sub>SB</sub> | 0.298 | 1.074 |
| *Carex acutiformis*<sub>NSB</sub> | 0.277 |  |
| *Cirsium arvense*<sub>SB</sub> | 0.196 | 1.226 |
| *Cirsium eriophorum*<sub>NSB</sub> | 0.160 |  |
| *Galium aparine*<sub>SB</sub> | 0.158 | 0.751 |
| *Galium odoratum*<sub>NSB</sub> | 0.210 |  |
| *Geranium pusillum*<sub>SB</sub> | 0.342 | 1.000 |
| *Geranium pyrenaicum*<sub>NSB</sub> | 0.342 |  |
| *Lamium album*<sub>SB</sub> | 0.171 | 0.903 |
| *Lamium galeobdolon*<sub>NSB</sub> | 0.189 |  |
| *Poa trivialis*<sub>SB</sub> | 0.128 | 1.059 |
| *Poa nemoralis*<sub>NSB</sub> | 0.121 |  |
| *Ranunculus flammula*<sub>SB</sub> | 0.157 | 0.888 |
| *Ranunculus reptans*<sub>NSB</sub> | 0.176 |  |
| *Silene vulgaris*<sub>SB</sub> | 0.183 | 1.030 |
| *Silene uniflora*<sub>NSB</sub> | 0.178 |  |

**Figure S1.**
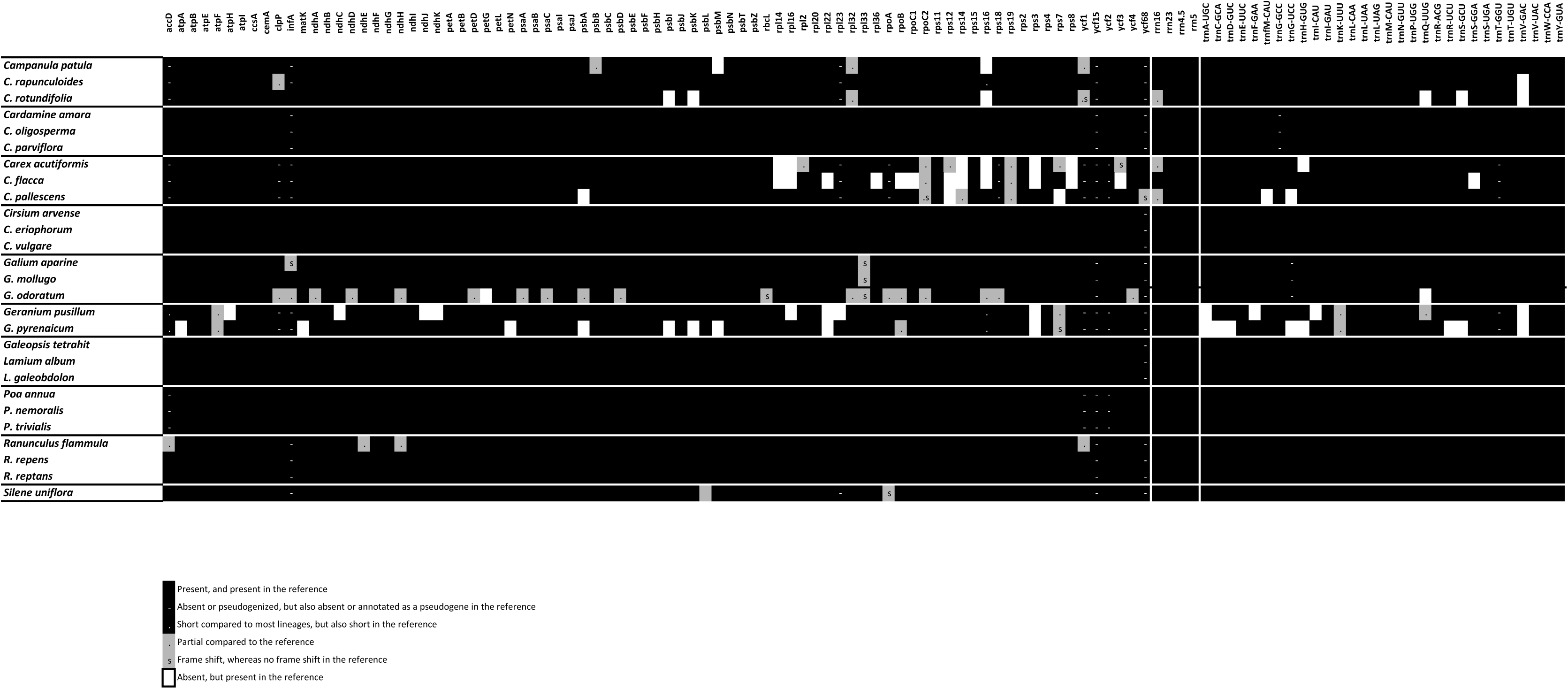
Plastid gene content for the new plastomes. White boxes indicate the absence of a gene that is present in closely related species. Grey boxes indicate genes that are partial (dots) or contain frameshifts (s) whereas they are present in closely related species as longer complete open reading frames. Black boxes indicate genes that show the same stage in the reference, with filled boxes indicating genes present and apparently functional, whereas minus signs indicate an absence or possible loss of function (always both in the taxon of interest and closely related species. Therefore, whereas white and grey boxes frame shift) of the gene, and dots a short size compared to other lineages, indicate features that could be artefacts due to low-coverage regions, black boxes indicate biological features.

**Figure S2.**
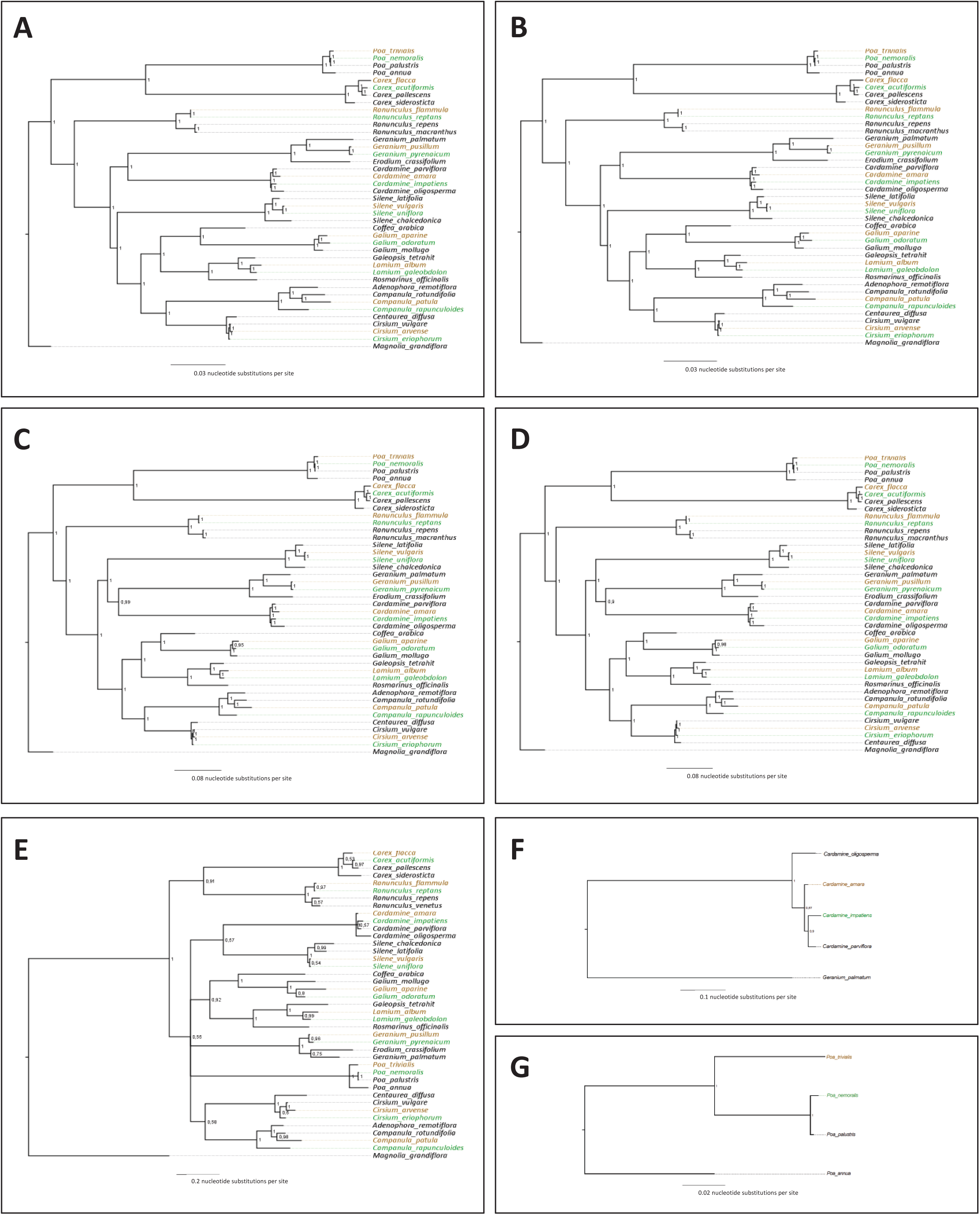
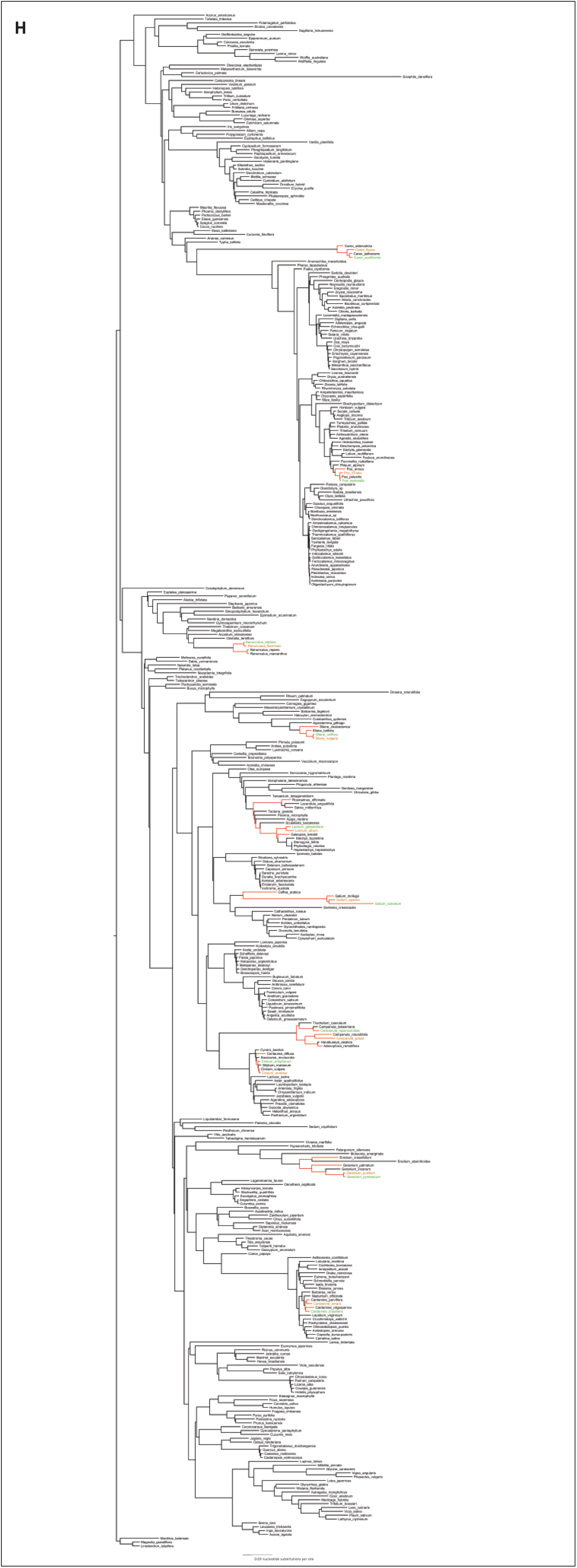
Bayesian phylogenetic trees used for relative substitution rate analyses. A. Topology obtained with the concatenated plastid coding sequences (CDS). B. Topology obtained with the CDS skimmed of loci yielding single-gene topologies conflicting the topology derived from all concatenated CDS. C. Topology obtained with the concatenated plastid non-coding sequences (NCS). D. Topology obtained with the NCS skimmed of loci yielding single-gene topologies conflicting the topology derived from all concatenated NCS. E. Topology obtained with the Nuclear ITS. F. Topology obtained with the nuclear ITS of *Cardamine*, with *Geranium palmatum* as outgroup. G. Topology obtained with the nuclear ITS of *Poa*. H. Topology obtained with the concatenated plastid CDS of 399 angiosperm taxa, including those used in this study. All topologies shown here are derived from non-partitioned datasets, but those from partitioned datasets are qualitatively identical (see main text and Tables S2 and S3).

**Figure S3.**
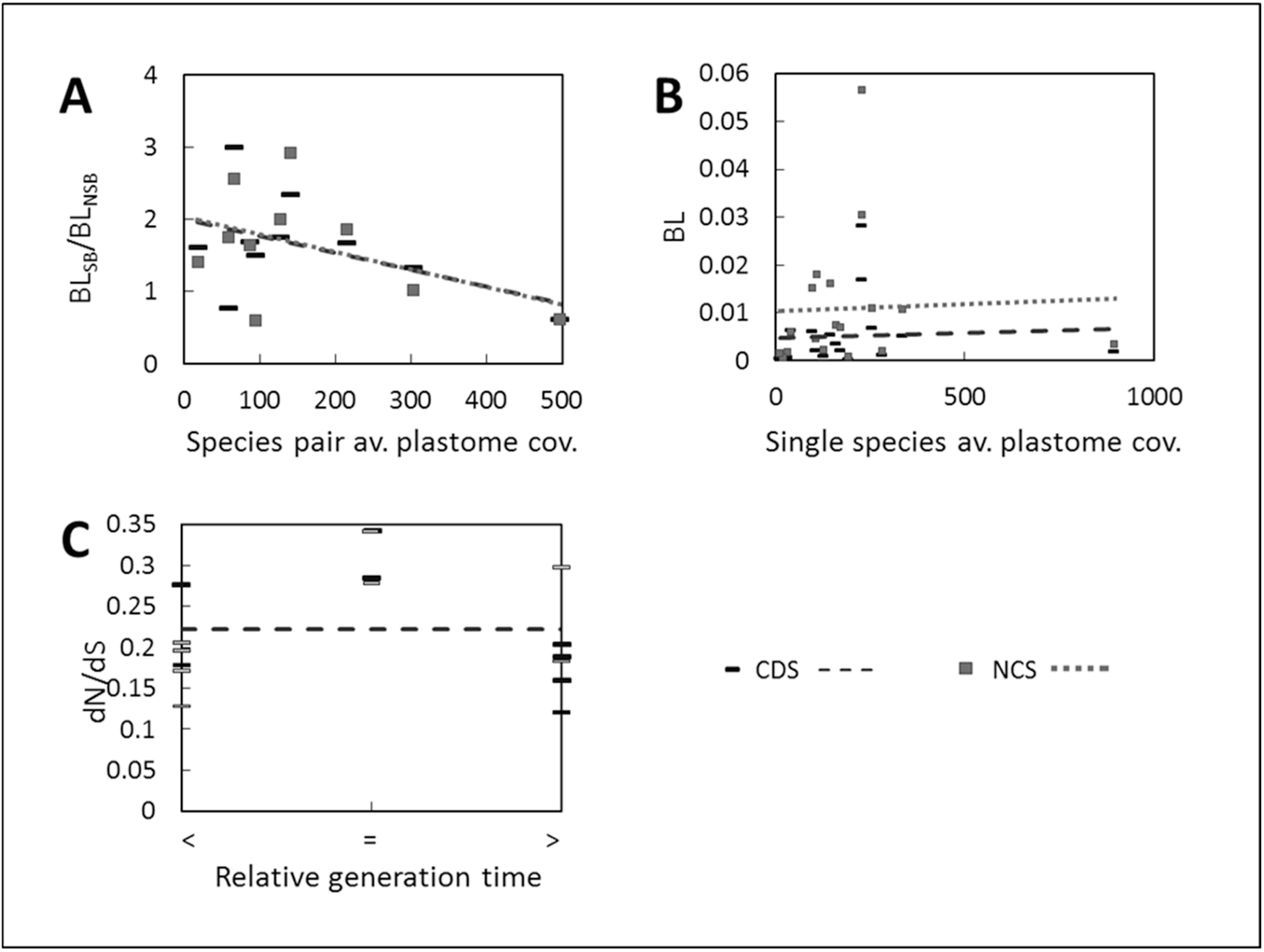
Correlation analyses of substitution rates as a function of additional factors. A. Branch-lengths ratios BL_SB_/BL_NSB_ (from the analysis of non-partitioned concatenated sequence alignments) as a function of average plastome coverage of the species pair. B. Branch-length as a function of average plastome coverage of individual species. C. dN/dS as a function of generation time (relative to that of the other species of the pair). In C grey bars indicate SB species and black bars indicate NSB species. The inferior, equal and superior to signs indicate respectively that the species had a smaller, equal or larger generation time than its sister species.

**Table S1. Species sampled in this study,** with their voucher information, and accession numbers of plastomes newly generated and previously published. Information on the starting material and DNA quality, as well as coverage statistics are also provided.

**Table S2. Branch lengths obtained from all Bayesian phylogenetic analyses.** Branch lengths were not estimated for the genus-level trees of *Cardamine* and *Campanula* due to the impossibility to root them outside the species pair of interest.

**Table S3. Summary of the substitution rate differences between seed-bankers and non-seed-bankers obtained in all Bayesian phylogenetic analyses.** Branch lengths were not estimated for the genus-level trees of *Cardamine* and *Campanula* due to the impossibility to root them outside the species pair of interest.

**Table S5. Biological information on the species involved in seed-banker/non-seed-banker comparisons.**

**Table S7. Accession numbers of whole plastomes, nuclear ITS and plastid regions used to build Figure S2H and to perform polymorphism analyses in *Lamium* and *Cardamine*.**

